# Tick-borne coinfections modulate CD8+ T cell response and progressive leishmaniosis

**DOI:** 10.1101/2025.04.03.647089

**Authors:** Breanna M. Scorza, Danielle Pessôa-Pereira, Felix Pabon-Rodriguez, Erin A. Beasley, Kurayi Mahachi, Arin D. Cox, Eric Kontowicz, Tyler Baccam, Geneva Wilson, Max C. Waugh, Shelbe Vollmer, Angela Toepp, Kavya Raju, Oge Chigbo, Jonah Elliff, Greta Becker, Karen I. Cyndari, Serena Tang, Grant Brown, Christine A. Petersen

## Abstract

*Leishmania infantum* causes human Visceral Leishmaniasis and Leishmaniosis (CanL) in reservoir host, dogs. As infection progresses to disease in both humans and dogs there is a shift from controlling, Type 1, immunity to a regulatory, exhausted, T cell phenotype. In endemic areas, association between tickborne coinfections (TBC) and *Leishmania* diagnosis and/or clinical severity has been demonstrated. To identify immune factors correlating with disease progression, we prospectively evaluated a cohort of *L. infantum* infected dogs from 2019-2022. The cohort was TBC-negative with asymptomatic leishmaniosis at the time of enrollment. We measured TBC serology, anti-*Leishmania* antigen T cell immunity, CanL serological response, parasitemia, and disease severity to probe how nascent TBC perturbs the immune state. At the conclusion, TBC+ dogs with CanL experienced greater increases in anti-*Leishmania* antibody reactivity and parasite burden compared to dogs that did not have incident TBC during the study. TBC+ dogs were twice as likely to experience moderate (LeishVet stage 2) or severe/terminal disease (LeishVet stage 3/4). Prolonged exposure to TBC was associated with a shift in *Leishmania* antigen-induced IFNγ/IL-10 and enhanced CD8 T cell proliferation. Frequency of proliferating CD8 T cells significantly correlated with parasitemia and antibody reactivity. TBC exacerbated parasite burden and immune exhaustion. These findings highlight the need for combined vector control efforts as prevention programs for dogs in *Leishmania* endemic areas to reduce transmission to humans. Public health education efforts should aim to increase awareness of the connection between TBC and leishmaniosis.

## INTRODUCTION

Disease caused by *Leishmania infantum*, Visceral Leishmaniasis (VL) in humans and Leishmaniosis (CanL) in the reservoir host, dogs, is fatal in both species if not treated (1,2). Only a fraction of infected human or canine hosts progress to disease for reasons still poorly understood (3,4). Host cell microbicidal mechanisms have been shown to prevent establishment of intracellular infection within phagocytic cells. During controlled infection, LeishVet stages 1 and 2, a Type 1 immune response dominated by IFNγ-secreting CD4 T cells contributes to maintaining low parasite burden (5–7). A shift toward a regulatory immune environment with concomitant immunosuppression diminishing the host’s ability to constrain parasite replication can lead to high IL-10 production, pathology-inducing levels of circulating antibodies (hypergammaglobulinemia), reduced T cell reactivity, parasite outgrowth, and pancytopenia (2,8). This manifests clinically as anemia, malaise, and cachexia, characteristic of VL, with the addition of renal disease in CanL as well as HIV-coinfected people with VL, marked by elevated creatinine and proteinuria, a serious complication associated with high mortality rates (2). The temporal changes and factors that drive the shift from controlling Type 1 immunity to a regulatory, exhausted T cell phenotype are poorly understood (7,9). Increasing parasite burden and moderate clinical disease were associated with increased transmission from dogs to sand fly vectors (10).

Therefore, understanding what immune factors correlate with disease progression using dogs, both as a model and a critical reservoir host, is crucial to interrupting the transmission cycle for overall public health.

VL is a neglected tropical disease found in regions where comorbid challenges predispose the host to progressive disease. Coinfection with a variety of pathogens from immunosuppressive viruses to immune-shifting helminth infections have been correlated with higher likelihood of VL (11,12). This phenomenon also occurs in the canine reservoir, where common coinfections are tickborne bacterial pathogens. A causal association between tickborne coinfections (TBCs) and rate of *Leishmania* positive diagnosis and/or clinical severity has been demonstrated in endemic areas (13). To better understand the immune alterations that occur in naturally infected hosts prior to disease progression, we followed a cohort of *L. infantum* infected dogs from 2019-2022 to perform a prospective evaluation of anti-*Leishmania* antigen immunity and how inflammatory alterations like comorbid tickborne infections perturb the immune state leading to progression with severe and/or fatal consequences.

We hypothesize that TBCs modulate the anti-*Leishmania* immune response to negatively impact parasitic control, pathogenic antibody progression, and disease. TBCs are acutely inflammatory (14), which may push a shift toward a regulatory environment in the setting of chronic *Leishmania* infection and promote onset of immune exhaustion. During this study we identified the kinetics and immune markers associated with this critical shift and resultant progressive leishmaniosis.

## MATERIALS AND METHODS

### Ethics statement

All procedures involving animals during this study were approved by the University of Iowa Institutional Animal Care and Use Committee and performed under the supervision of licensed and, where appropriate, board-certified veterinarians according to International AAALAC accreditation standards.

### Animals/cohort selection

We identified a cohort of naturally *L. infantum*-exposed dogs without clinical CanL and seronegative for the most common endemic canine tickborne pathogens: *Borrelia burgdorferi, Anaplasma phagocytophilum, Anaplasma platys, Ehrlichia chaffeensis, Ehrlichia canis,* and *Ehrlichia ewingii*. This cohort was selected from a population of client-owned hunting hounds in which *L. infantum* is enzootic across three sites in the Midwestern U.S.

(15). Due to the working nature of this cohort, there is relatively high exposure to ticks and tickborne pathogens compared to companion dogs, similar to dogs in endemic areas of Brazil or Southern Europe (15). This cohort provided us a unique opportunity to monitor immune variables over time, as dogs are naturally exposed to environmentally occurring TBCs and concomitant CanL development.

Our research group assayed cohort *Leishmania* and TBC diagnostics, staged clinical CanL, and performed *ex vivo* T cell restimulation assays in response to *Leishmania* antigen at three-month intervals over a course of 18 months, inclusive of two tick seasons, with additional clinical follow-up at 2-and 2.5-years post-enrollment. U.S. hunting dogs from a naturally CanL enzootic cohort were screened for *Borrelia burgdorferi*, *Anaplasma* spp., and *Ehrlichia* spp. serology via IDEXX 4Dx SNAP test (IDEXX Reference Labs Inc.) and *Leishmania* serology via CVL®DPP. DNA was isolated from whole blood and analyzed for presence of *Leishmania* DNA via RT-qPCR described below. Physical exam was performed by licensed veterinarians for clinical signs of leishmaniosis (16).

Fifty dogs were enrolled based on inclusion criteria: positive *Leishmania* diagnostic test (DPP or qPCR) or *Leishmania* diagnostic positive dam or full sibling (DPP or qPCR); seronegative 4Dx SNAP test; two or fewer physical clinical signs of CanL. Dogs were excluded if they met any of the following criteria: negative *Leishmania* diagnostic test results and no *Leishmania* diagnostic positive dam or full sibling; CVL®DPP serology value > 200; three or more physical clinical signs of leishmaniosis; positive 4Dx SNAP test. Previous work in the same enzootic population identified that dogs born to dams ever positive for *Leishmania* had a 13.8-fold greater risk of becoming diagnostically positive themselves (p=0.0002) (21).

### Study design

Cohort immune responses were followed over 18 months from April 2019 to November 2020 (Supplemental Figure 1). At three-month time points (Figure 1), whole blood and serum were collected and physical exam performed to clinically stage disease. *L. infantum* serology, *L. infantum* blood parasite burden, and *Borrelia burgdorferi*, *Anaplasma* spp., and *Ehrlichia* spp. serology via IDEXX 4Dx SNAP test (IDEXX Reference Labs Inc.) were performed. After the conclusion of the main study period, remaining subjects were visited in the following year (2021) by the research team to evaluate progression and mortality outcomes. At the time of final analysis for this manuscript, dogs that never had positive PCR or serological *Leishmania* diagnostic result were censored from analyses (n=9).

**Figure 1.**
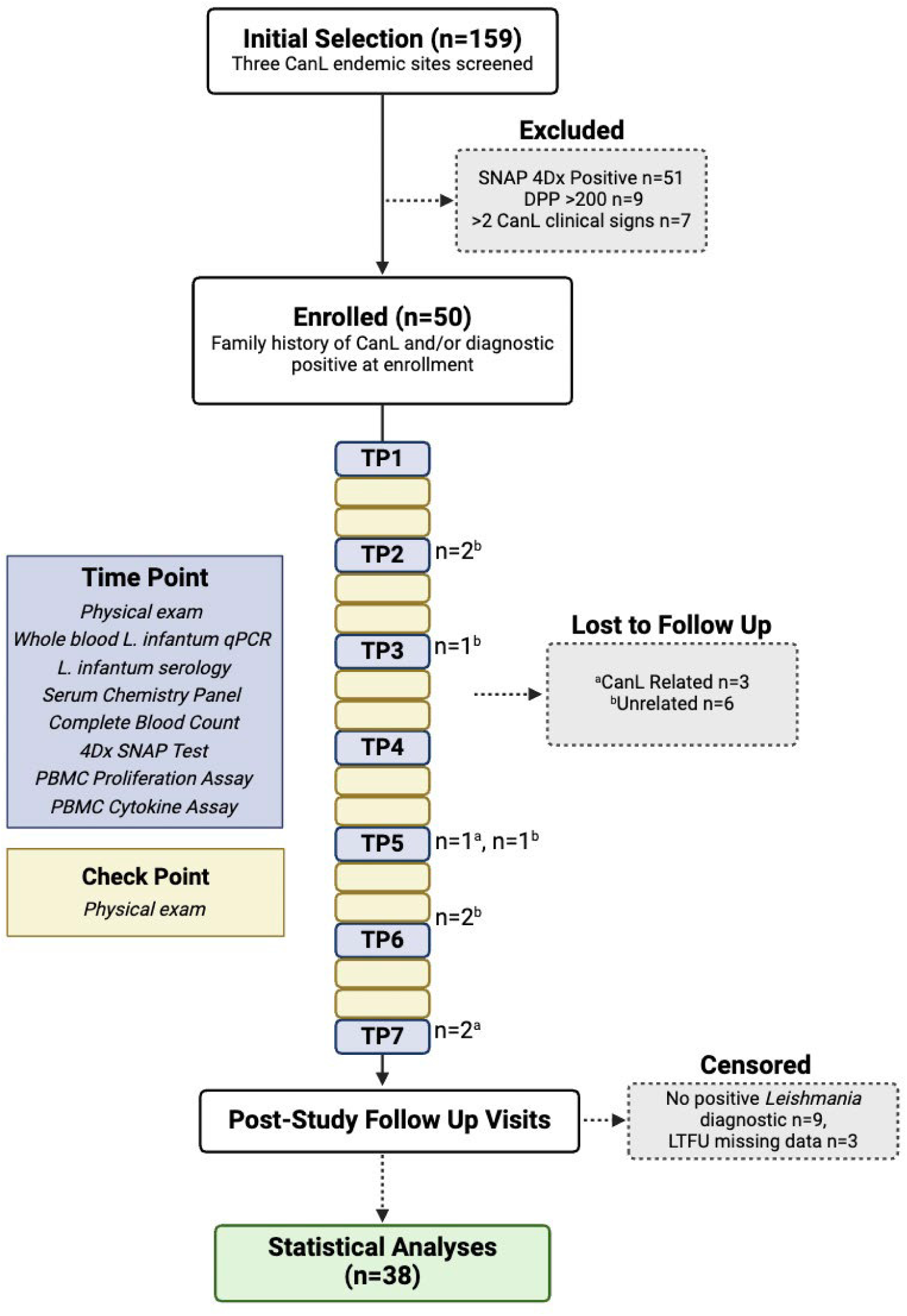
Prospective study fllow chart. A cohort of 50 dogs with asymptomatic *Leishmania infantum* exposure and seronegative for tick-borne pathogen exposure via 4Dx SNAP test were followed for 18 months. Evaluations performed at each major time point (TP) occurring every three months, check point (CP) occurring every month, and number of dogs lost to follow up at each site visit provided.

To control for highly inflammatory bacterial TBC on immunity, dogs were block randomized by sex and age into two treatment groups receiving either oral tick prevention (Sarolaner, Zoetis Inc.) or placebo. Treatment was administered monthly in a double-blinded manner; caretakers, veterinarians, and researchers were blinded to treatment status throughout the study.

### Clinical staging

Clinical leishmaniasis was staged according to the LeishVet staging system (2). Venous blood was drawn at three-month intervals. Complete blood count and serum chemistry panels were performed by IDEXX Reference Laboratories (Westbrook, ME). Physical exam findings, complete blood counts, and serum chemistry panel results were systematically used to assign LeishVet scores (1–4). Dogs without any physical signs of disease or clinicopathological abnormalities were assigned as stage of 0 or “healthy.”

### Parasites and standard curve

For preparations of total *Leishmania* antigen and standard curves, *Leishmania infantum* promastigotes (US/2016/MON1/FOXYMO4), originally isolated from a hunting dog with CanL, were cultured in complete hemoflagellate-modified minimal essential medium (HOMEM) with 10% heat-inactivated fetal calf serum at 26°C (17). Stationary phase parasites were enumerated on a hemacytometer and adjusted to 1×10^8^ parasites/mL in PBS. Ten-fold serial dilutions were made and 10 uL of each dilution was spiked into 1 mL of *Leishmania* negative control canine blood to create an exponential regression standard curve where circulating parasite load is calculated as parasite equivalents/mL canine blood (Supplemental Figure 2).

### Quantitative PCR

DNA was isolated from 200 µL of canine peripheral blood at each major study timepoint with the Gentra Puregene Tissue Kit (Qiagen). For *L. infantum* detection by Real Time-qPCR, *Leishmania* small-subunit rRNA-specific probe (5’-[6-FAM]-CGGTTCGGTGTGTGGCGCC-MGBNFQ-3’) and primers (forward, 5’-AAGTGCTTTCCCATCGCAACT-3’; reverse, 5’-GACGCACTAAACCCCTCCAA-3’) were used as previously described (18).

### Preparation of Leishmania antigen

*Leishmania* antigen was prepared after washing stationary phase *L. infantum* promastigotes in cold PBS twice followed by multiple freeze-thaws in 5 mM CaCl_2_, 10 mM Tris, pH 7.4. Lysed parasite solution was either used as is (Total *Leishmania* Antigen, TLA) or centrifuged at 5000 RCF for 20 minutes at 4°C and supernatant collected (Soluble *Leishmania* Antigen, SLA). Protein content of lysate or supernatant was quantified via Pierce BCA protein assay (Thermo Scientific) and aliquots frozen at-30°C.

### Leishmania ELISA

For indirect ELISA, 96-well flat bottom EIA treated plates were coated with 2 µg/mL SLA in bicarbonate buffer (pH 9.6) overnight, blocked with 1% w/v bovine serum albumin + 0.05% w/v Tween-20 in PBS, and probed with canine sera diluted 1:500 in blocking buffer. All samples and controls were run in duplicate. Peroxidase-conjugated AffiniPure anti-dog IgG (H+L) (Jackson ImmunoResearch) was diluted 1:20,000 in blocking buffer, and OptEIA TMB substrate (BD Biosciences) was used to detect peroxidase activity. The reaction was stopped with 2M H_2_SO_4._ Absorbance was read at 450 nm and 650 nm on a spectrophotometer plate reader.

Serum samples from five CanL-negative dogs, from a non-endemic area, were included on each plate. A cut-off was calculated from the average absorbance of the negative controls plus three standard deviations. Test serum samples were considered positive if their average absorbance was greater than the cut-off. SLA ELISA results are reported as test sample absorbance ratio over the control cut-off. An SLA ELISA ratio >1 is considered positive.

### PBMC stimulation assay

Peripheral blood mononuclear cells (PBMCs) were isolated from whole blood via density centrifugation over Ficoll-Paque PLUS (Cytiva, Fisherbrand). PBMCs were plated in 96-well round bottom plates at 5×10^5^ cells/well in RP-10 media (RPMI 1640 supplemented with 10% FBS, 2 mM *L*-glutamine, penicillin-streptomycin, and non-essential amino acids). A subset of PBMCs were stained with CFSE prior to stimulation. CFSE labeling was performed in the dark. CFSE was prepared at 5 µM in PBS. PBMCs at 5×10^6^ cells/mL were mixed with an equal volume of CFSE solution (final staining concentration 2.5 µM), incubated at room temperature for 5 minutes, and quenched with RP-10 for 5 minutes at room temperature.

PBMCs were stimulated with media, 10 µg/mL TLA, 1 µL/well canine distemper virus vaccine antigen (DHPP-a relevant antigen positive control), or 5 µg/mL Concanavalin A (ConA). Supernatants were collected from PBMCs receiving each stimulus after 72 hours. Flow cytometry was performed on PBMCs stimulated with media, TLA, or DHPP for 7 days and with ConA for 3 days. For intracellular cytokine staining, 10 µg/ml brefeldin A (Sigma-Aldrich) was added 6 hours prior to harvest.

### Cytokine ELISAs

Supernatants from PBMCs restimulated with media, TLA, DHPP, or ConA as described above were collected after 72 hrs. Sandwich ELISA was performed on supernatants using canine specific IFNγ, IL-10, and TNF DuoSet ELISA kits according to the manufacturer instructions (R&D Systems).

### Flow cytometry

Cells were blocked with 50% FBS/PBS containing 0.9 µL/well dog gamma globulin for 15 min (Jackson ImmunoResearch). Surface labeling antibodies were prepared in block solution: anti-canine CD4-AF647 (1:250, BioRad antibodies), anti-canine CD8-AF700 (1:250, BioRad antibodies), biotinylated anti-human PD-1 (1:40, R&D) and anti-human CD49d-BV605 (1:50, BioLegend Inc). Streptavidin-PE-Cy7 secondary label was used according to manufacturer instructions (BioLegend Inc). Cells were labeled on ice in the dark for 30 min. Samples were fixed for 15 min in fixation buffer (BioLegend Inc), washed with PBS, and stored at 4°C protected from light.

For intracellular cytokine staining (ICS), fixed cells were permeabilized in 1X perm/wash solution for 15 min room temperature in the dark. Zenon conjugation kits were used to label anti-canine IFNγ-R-PE (5 µg/mL, R&D Systems) and anti-canine IL-10-AF488 (5 µg/mL, R&D Systems) antibodies (ThermoFisher Z25255 and Z25002, respectively) according to manufacturer instructions. Cells were labeled on ice for 30 minutes in the dark. Cell events were acquired within 48 hours on a Becton Dickenson LSR II flow cytometer and data were analyzed using FlowJo software (Supplemental Figures 3 and 4).

### Statistical analyses

Longitudinal modeling is described in the following section. All other statistical analyses were performed using GraphPad Prism software Version 10.0.3. Statistical tests applied can be found in the figure legend for each comparison. Normality tests were applied to determine if data should be analyzed with parametric or nonparametric test. If any normality test indicated non-normality, the test was performed as nonparametric.

Unless indicated, outliers were not removed from analyses. Where indicated in figure legend, outliers were identified and removed using the ROUT test.

### Bayesian linear mixed effects model

To better understand the temporal changes of the overall immune response, comorbid TBC, and clinical outcome by the different immune measurements, we used a Bayesian linear mixed effects model. The timepoint of TBC seroconversion of each TBC+ study subject was denoted *t_S_*. The analysis included six study time points for each TBC+ subject, including *t_s_*, three and six months prior to *t_S_*, and three, six, and nine-months post *t_S_*. The explanatory variables were normalized to mean zero and variance 1. This allows the inclusion of explanatory variables at multiple scales, while keeping the interpretation of the magnitude of regression coefficients comparable. LeishVet status (ordinal) was treated as a numeric variable for clinical outcome to incorporate into the model, so effects are interpreted in terms of their linear effect on changes in LeishVet score. Bayesian imputation (19) was used to impute unobserved values in the dataset.

Given *Y_ij_* represents a study variable of interest for subject *i* at time j, individual immunopathogenic variables were modeled using the following specification:

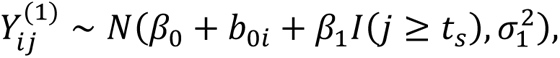

where β_0_ denotes the overall intercept (fixed effect), *b_0i_* the subject-specific random intercept, β_1_ the effect of TBC+ time on *Y*(^1^)_ij_, and ϵ_ij_ the error terms such that ϵ_ij_ ∼ N(0, σ_b_^2^) and *b_0i_* ∼ N(0, σ_b_^2^). σ_1_^2^ and σ_b_^2^ denote population and individual-specific variances, using inverse-gamma prior distributions. β_0_ and β_1_ used normal prior distributions. The mean structure included the indicator function I(j ≥ *t_s_*), which assigned a value of 1 at timepoints including or following *t_S_*, and 0 at timepoints preceding *t_S_*.

Three Markov Chain Monte Carlo simulations of 25000 iterations and 5000 iterations as a burn-in period were run, for a total of 20000 iterations on each chain. To assess parameter convergence, we used the Gelman-Rubin diagnostic from the coda package (20) in R. All parameters in the model reached convergence.

With this model specification, we are interested in the posterior distribution of β_1_, the impact of comorbid infection on clinical status, and negative or positive direction of the posterior probability. We evaluate the strength of the evidence for an association between comorbid infection and clinical status through max(P(β_1_ > 0| ⋅), P(β_1_ < 0| ⋅)), the higher one-tail probability. This probability is the criterion used to assess the strength of evidence for the corresponding hypothesis that TBC modulates each study variable being modeled. We considered the posterior probability (PP) of the TBC effect (positive or negative) on a given variable of interest as moderate, if the PP lies in the interval [0.65, 0.85], while a PP in the interval [0.85, 1] was considered a strong effect. Supplemental Table 1 shows all PP results.

## RESULTS

### Cohort inclusion and study design

For this prospective CanL cohort study, 159 subjects were screened and excluded based on listed exclusion criteria (Figure 1). This excluded subjects with clinically apparent CanL and subjects exposed to current or recent TBC. Of 92 candidates, 50 dogs were enrolled in the study; 24 dogs with positive *Leishmania* diagnostic testing confirmed through serological or molecular testing, and 26 dogs that had either a dam or full sibling with a positive *Leishmania* diagnosis. Together, these inclusion and exclusion criteria resulted in a cohort of dogs with confirmed or highly suspected *in utero Leishmania* exposure but not presenting clinical CanL or comorbid TBC at the time of enrollment.

The enrolled cohort was followed over the course of 18 months between 2019-2020, with physical examination by Petersen Lab veterinarians monthly and blood collection for immunoassays and diagnostics every three months (Figure 1, Supplemental Figure 1A). This cohort was comprised of 52% male and 48% female dogs, 42% aged 0-2 years, 48% aged 3-5 years, and 10% aged 6 years or greater (Supplemental Figure 1C).

Throughout the course of the study, nine subjects were lost to follow-up (LTFU) and had partial missing data for CanL-related (n=3) or non-CanL related circumstances such as ran away, not present on location during site visit, and behavioral issues (n=6) (Figure 1). This loss was determined to be at random.

After the study period concluded, remaining subjects were assessed for long-term CanL outcomes by physical examination, diagnostic testing, and caretaker-provided health history for 0-19 months. Thirty-eight dogs had a positive *Leishmania* diagnosis at the end of the trial and follow-up monitoring period. Twelve subjects were censored from data analysis due to never testing positive on a *Leishmania* diagnostic test (n=9) or were LTFU and did not have sufficient data (n=3). Therefore, data from a total of 38 confirmed *Leishmania*-positive canine subjects were included in the analyses presented (Figure 1).

### Tickborne coinfections exacerbate clinical presentation and CanL severity

At the end of the study, 19 dogs tested TBC-positive on the IDEXX 4Dx SNAP test at some point during the study period (Ever TBC+). We wanted to establish the extent to which TBC impacted CanL serological response and parasitemia, as measured by ELISA, qPCR and disease severity. When comparing subject-matched first and final measurements for each group, we observed that both TBC-unexposed (p=0.008) and TBC-exposed (p<0.0001) CanL dogs experienced a significant increase in blood parasite burden across the study period, demonstrating natural progression of CanL (Figure 2A). Anti-*Leishmania* serological reactivity also increased significantly in both TBC unexposed (p=0.012) and TBC exposed (p<0.0001) dogs across the study period (Figure 2C). At the study conclusion, a larger proportion of TBC+ dogs had become parasitemic or seropositive for *Leishmania* compared to TBC-dog (Figure 2B and D), but this difference was not statistically significant.

**Figure 2.**
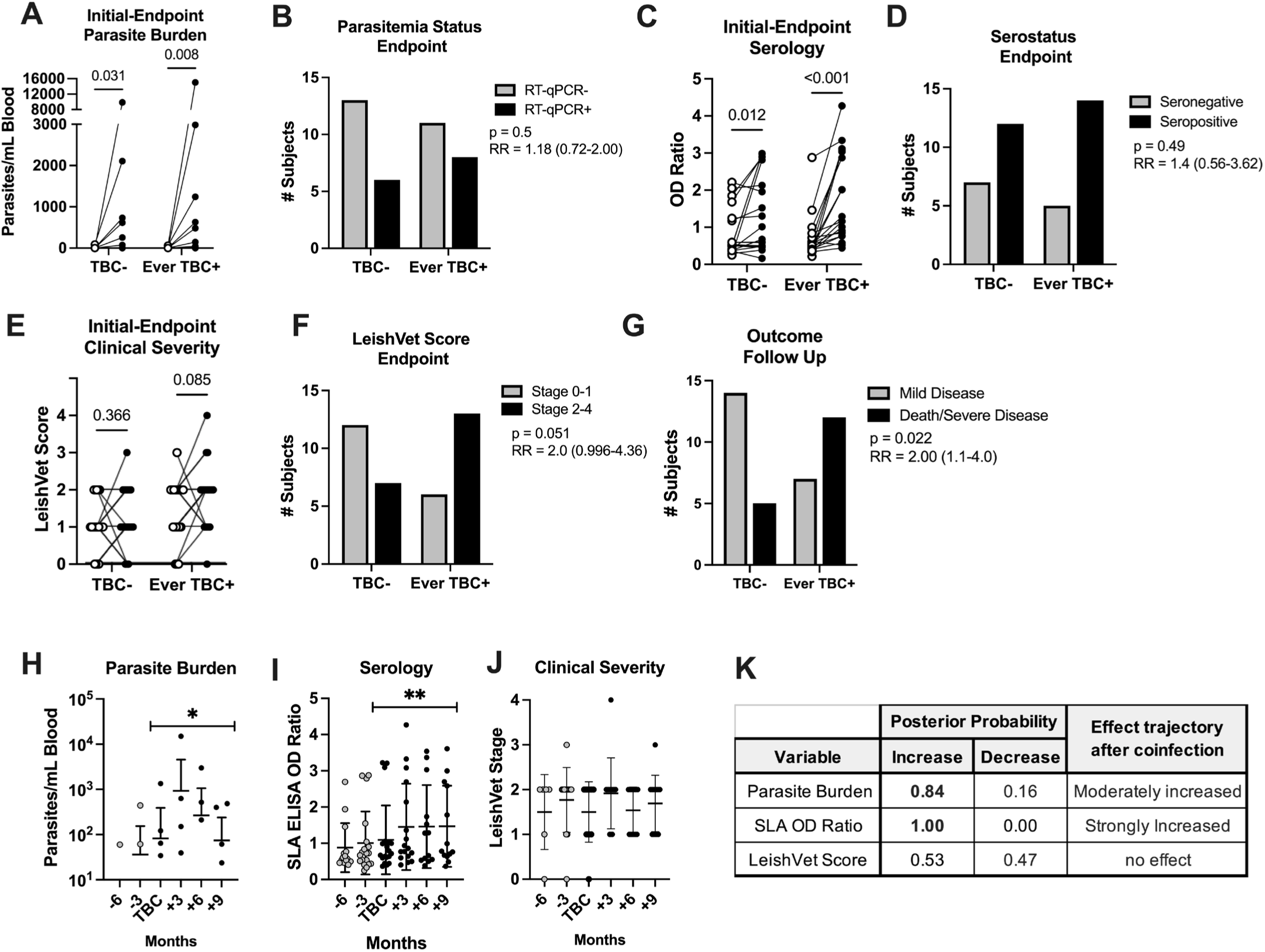
**Significant progression of CanL in dogs with TBC exposure**. *L. infantum* qPCR (A), SLA OD Ratio (C), and LeishVet score (E) of TBC-or TBC+ subjects. Wilcoxon matched-pair signed rank test. A, C, E. Subject matched initial timepoint value (white) compared to endpoint value (black) of blood parasite burden (B), serological status (D), LeishVet score (F), or clinical outcome at post-study follow up timepoint (G) between TBC-or ever TBC+ subjects. Chi-squared contingency analysis of parasitemia status by RT-qPCR Chi-squared test p-value and relative Risk (RR) with 95% confidence interval. TBC-, subjects that never experienced a TBC. Ever TBC+, subjects that were TBC+ at any point during the study. (H-K) Kinetics of *Leishmania* diagnostics among TBC-positive subjects at three-month intervals preceding and following timepoint of TBC seroconversion. Posterior probability (PP) of change in behavior after TBC seroconversion is indicated. *PP 0.65-0.85, moderate effect. **PP ≥ 0.85, strong effect. Median with 95% CI shown.

Using the LeishVet clinical severity scale, dogs with subclinical CanL exposed to a TBC experienced a greater increase in LeishVet scores across the study period as compared to those unexposed (Figure 2E). At their final study assessment, TBC+ dogs were at twice the risk of being assigned a LeishVet score of 2 or higher compared to TBC-dogs (relative risk=2.0, p=0.051) (Figure 2F). Further, at follow-up visits, dogs that experienced a TBC during the study period (TBC+ dogs) had double the risk of developing severe disease or undergoing euthanasia due to leishmaniosis. Dogs that remained TBC-during the study period were more likely to remain subclinical or maintain mild disease (p=0.022, relative risk= 2.0) (Figure 2G). Together, these data support the conclusion that dogs with subclinical CanL experiencing a TBC during the study period significantly increased the likelihood of progression to clinical CanL by the completion of the study.

Using Bayesian linear mixed-effects modeling, we focused specifically on TBC+ dogs within our cohort and modeled how clinical diagnostic measurements varied over time, considering the first incidence of TBC+ test result as an inflection point and comparing the change in variable behavior prior to coinfection versus after the tick pathogen exposure event. The resulting posterior probability (PP) of the model indicates the direction (increase, decrease, or no change) of the behavior kinetics across the coinfection timepoint and the magnitude of the probability of this behavior trend compared to pre-coinfection timepoints.

Applying this statistical model to TBC+ dog data, parasite burden and anti-*Leishmania* serological reactivity showed a moderate to strong increasing trend following the TBC event compared to the six months preceding TBC (Figure 2H-K). This supports TBC as a triggering event for observed increased parasitemia and serological OD.

### Tick-borne coinfections alter Leishmania antigen induced cytokine profile

Control of leishmaniasis is mediated by CD4^+^ T cells producing IFNγ to augment microbicidal activity of parasitized phagocytes (5–7). To counter this, *Leishmania* has evolved to induce numerous regulatory pathways that hamper pro-inflammatory responses. IL-10 induction during leishmaniosis is well documented *in vitro* and *in vivo* (8,22). IL-10 antagonizes the effects of IFNγ and other inflammatory cytokines, dampening damage from IFNγ, but chronic antigen exposure leads to high levels of IL-10, antibody production and high parasite burden (23). To assay cytokine production in our cohort, PBMCs from TBC+ CanL dogs were restimulated with *Leishmania* antigen (TLA) *ex vivo*.

IFNγ and IL-10 concentration was measured in supernatants by ELISA (Figure 3A and B) and cytokine-producing CD4^+^CD49d^hi^ lymphocytes were detected by intracellular staining and flow cytometry (Figure 3C and D).

**Figure 3.**
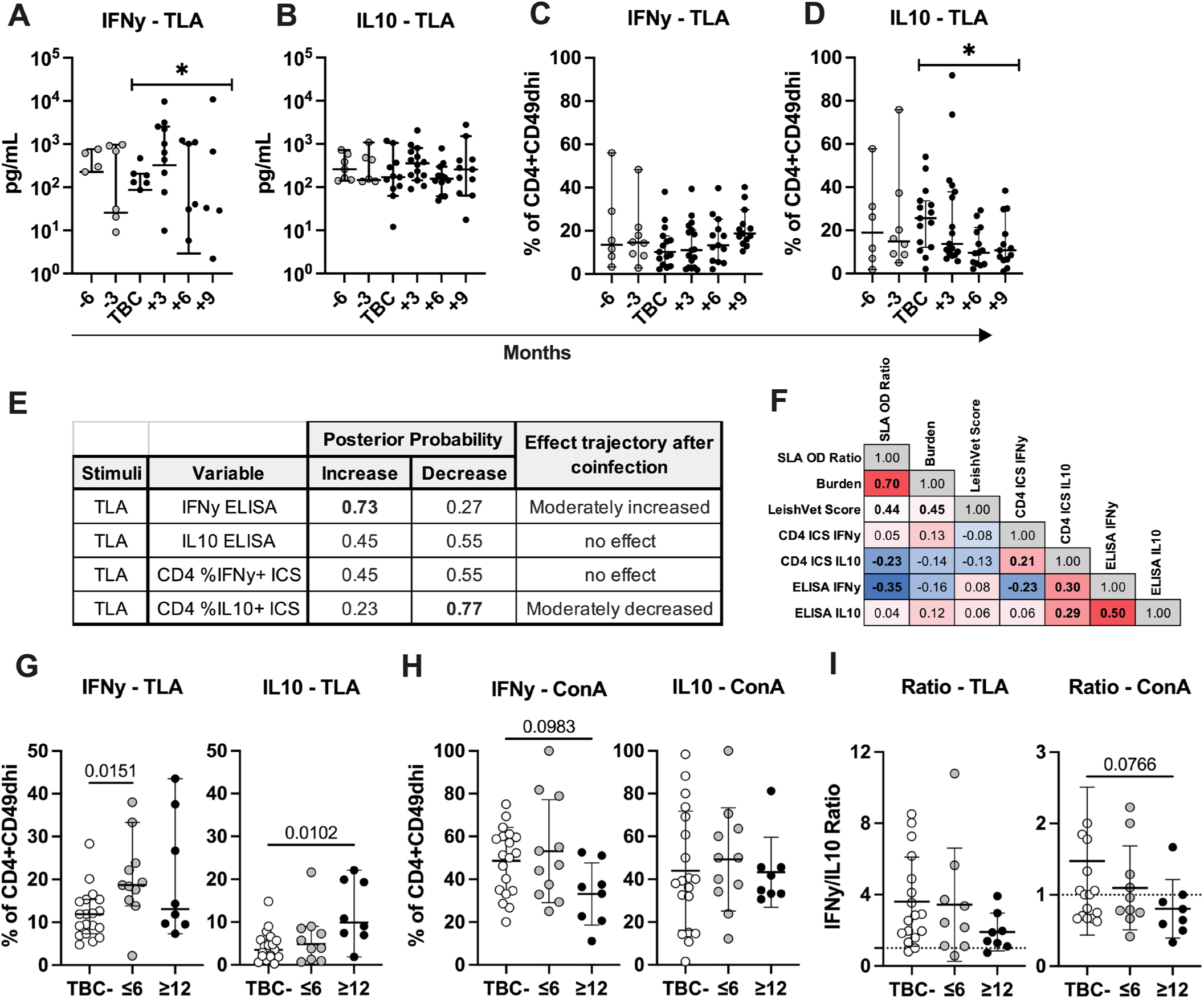
***Leishmania* antigen-stimulated IFN**γ **and IL-10 production following TBC.** Kinetics of TLA stimulated IFNγ (A,C) and IL-10 (B,D) production among TBC+ dogs at three-month intervals preceding and following TBC seroconversion. (A, B) cytokine concentration from stimulated PBMC supernatant via ELISA. (C, D) Frequency of cytokine expressing cells among TLA stimulated CD4^+^CD49d^hi^ lymphocytes by ICS. (E) Posterior probability of change after TBC seroconversion indicated. *PP 0.65-0.85, moderate effect. **PP ≥ 0.85, strong effect. Median with 95% CI shown. (F) Spearman correlation coefficient for each combination. Bolded values indicate p≤0.05. Heatmap shows red for strongest positive correlation coefficients and blue for strongest negative correlation coefficients. (G-I) Percent IFNγ or IL-10 positive CD4^+^CD49d^hi^ lymphocytes or ratio from TLA-or ConA-stimulated intracellular staining results at endpoint measurement of TBC-dogs, dogs TBC+ for ≤6 months or ≥12 months. Kruskal-Wallis with Dunn’s test or one-way ANOVA with Holm-Sidak test. Mean and SD provided.

Within TBC+ dogs, we observed a moderate increase in IFNγ secreted by bulk PBMCs in response to TLA following the TBC conversion event. However, the median IFNγ supernatant concentration became more variable with mean amounts trending downwards after TBC conversion (Figure 3A and E). IFNγ produced by TLA*-*stimulated PBMCs negatively correlated with parasite burden among TBC+ dogs (Figure 3F). TLA-induced IL-10 levels in bulk PBMC supernatants remained relatively steady up to nine months post-TBC (Figure 3B).

CD49d integrin expression differentiates antigen-experienced T lymphocytes from naïve cells (24). The frequency of IFNγ-producing CD4^+^CD49d^hi^ T cells did not significantly alter from the months prior to coinfection to post-TBC (Figure 3C and E). The proportion of IL-10^+^ CD4^+^CD49d^hi^ T cells moderately declined following the co-exposure event (Figure 3D and E) and significantly negatively correlated with circulating parasite burden (Figure 3F).

Following TBC seroconversion, dogs remained seropositive to tick pathogen antigen for variable periods of time. Of the 19 dogs that experienced TBC, 11 were seropositive against coinfecting species for 6 months or less, and 8 were seropositive for 12 or more months. Extended seropositivity could indicate ongoing coinfection or re-exposure. We hypothesized a prolonged coinfected state would have a more significant impact on the anti-*Leishmania* immune response and thus wanted to examine cytokine expression after shorter versus more chronic coinfection periods. To investigate this, we assessed cytokine production from TBC-, ≤6 months TBC+, and ≥12 months TBC+ dog cells (Figure 3G-J). To better visualize patterns between these groups, outliers were removed from comparisons of TBC duration.

At endpoint measurements, in response to TLA, ≤6 months TBC dogs showed a significant increase in % IFNγ^+^ CD4^+^CD49d^hi^ T cells compared to TBC-dogs (Figure 3G). However, dogs TBC+ for long-term periods (≥12 months), CD4^+^CD49d^hi^ T cells showed a significant increase in %IL-10 production (Figure 3G). In response to the mitogen ConA, we observed a trend of decreased IFNγ producing cells among long term TBC+ dog CD4^+^CD49d^hi^ T cells (Figure 3H).

The ratio of %IFNγ^+^/%IL-10^+^ CD4^+^CD49d^hi^ lymphocytes among long term TBC+ dog cells decreased non-significantly compared to TBC-dog cells at endpoint measurements (∼1.9-fold reduction in response to TLA and ∼1.8-fold reduction in response to ConA) (Figure 3I). Together, this data demonstrates a potential shift to a more regulatory profile in dogs co-exposed to *L. infantum* and TBC for extended time periods.

### Relationship between PD-1 expression and proliferation kinetics among coinfected dogs

In addition to active immune evasion by *Leishmania* parasites, T cell exhaustion can occur in settings of chronic antigen exposure and T cell activation such as visceral leishmaniasis, leading T cells to become less responsive to stimuli (25). Therefore, we measured surface Programmed Death 1 (PD-1) expression, a key marker of T cell activation but also exhaustion (26) on CD4^+^ and CD8^+^ T cells in relation to proliferation in response to *Leishmania* antigen. In dogs with CanL, TLA-stimulated CD4^+^CD49d^hi^ T cell proliferation (%CFSE^lo^) showed a strong probability of decreasing following the tick pathogen seroconversion timepoint (Figure 4A and E) and positively correlated with CD4^+^CD49d^hi^ T cell PD-1 expression across timepoints following TBC (r=0.73, p<0.001, Figure 4F). Looking more specifically at endpoint measurements, this correlation remained strongly significant in all dog groups TBC-, ≤6 mo TBC+, and long-term TBC+ (Figure 4I-K). Our results show that following TBC in CanL dogs, CD4^+^CD49d^hi^ cell proliferation decreases in a subject-matched, longitudinal model; however, PD-1 expression by these cells was highly correlated and thus does not explain this decrease, indicating other regulatory mechanisms are contributing to this decreased proliferative phenotype.

**Figure 4.**
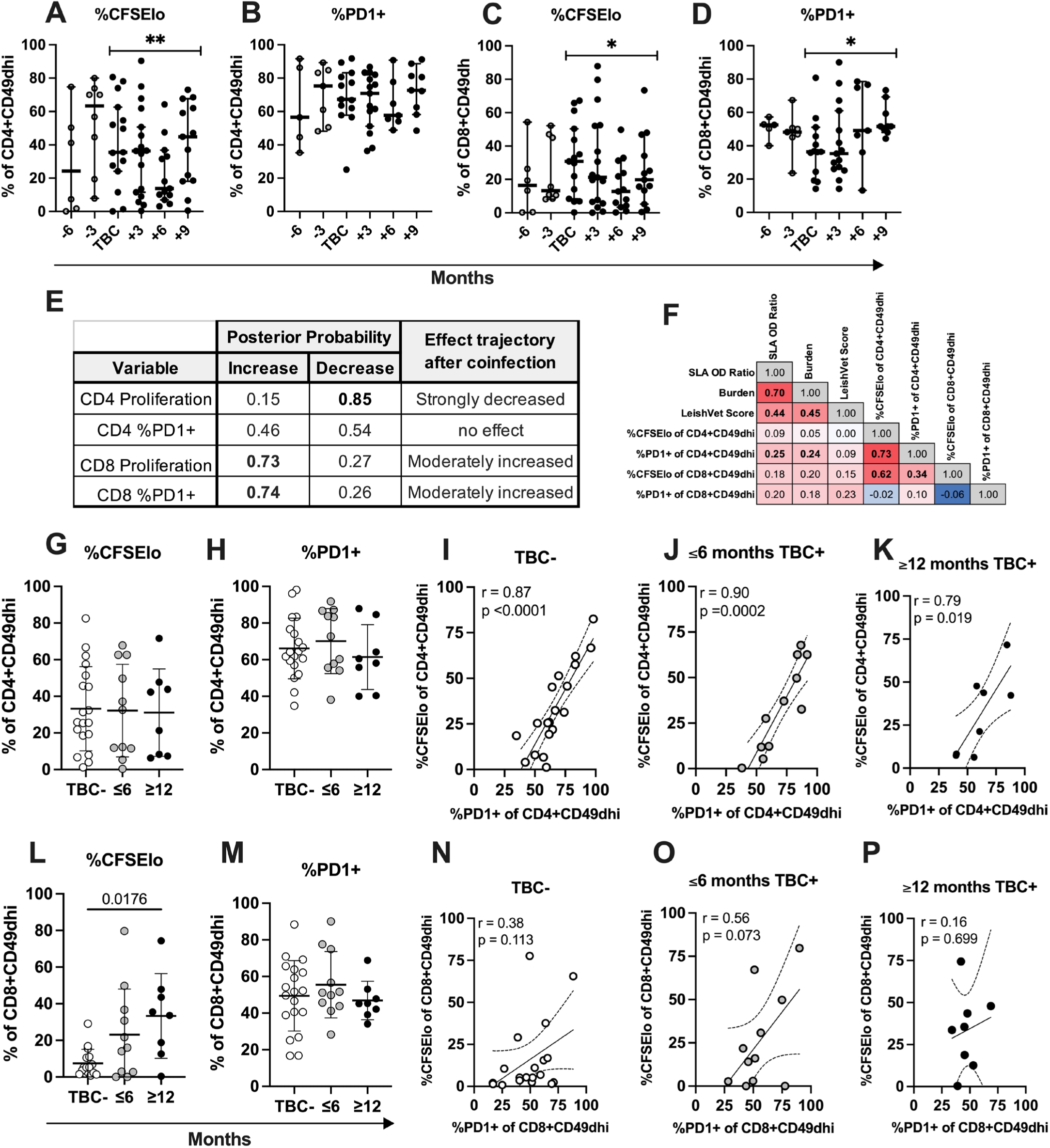
*Leishmania*-antigen stimulated CD4+ T cell proliferation and PD1 expression decreased after TBC while markers of CD8^+^ T cell activation increase. (A-E) Kinetics of TLA stimulated proliferation and PD1 expression as percent of CD4^+^CD49d^hi^ (A-B) or CD8^+^CD49d^hi^ (C-D) lymphocytes among TBC+ dogs at three-month intervals preceding and following timepoint of TBC seroconversion. Median and 95% CI shown. Posterior probability (PP) of change in behavior after TBC seroconversion is indicated. *PP 0.65-0.85, moderate effect. **PP ≥ 0.85, strong effect. (E) PP overview table. (F) Spearman correlation matrix of TBC+ dogs including all post-TBC measurements. Bolded values indicated p<0.05. Correlation coefficient shown. Heatmap shows red for strongest positive correlation coefficients and blue for strongest negative correlation coefficients. (G-H and L-M) Endpoint TLA stimulated proliferation and PD1 expression as percent of CD4^+^CD49d^hi^ (G-H) or CD8^+^CD49d^hi^ (L-M) lymphocytes among TBC-, dogs TBC+ for ≤6 months, or ≥12 months. Outliers removed. Mean and SD shown. One-way ANOVA with Hold-Sidak post-test. (I-K, N-P) Correlation between TLA stimulated proliferation and PD1 expression as percent of CD4^+^CD49d^hi^ (I-K) or CD8^+^CD49d^hi^ (N-P) lymphocytes among TBC-dogs or dogs TBC+. Spearman correlation coefficient (r) and p-value are shown. Linear regression and 95% CI are depicted.

Previous evaluation of CD8^+^ T cell responses to chronic *Leishmania* infection and disease indicated that these cells appeared to have an exhausted phenotype earlier in clinical progression compared to CD4^+^ T cells (7). How TBC and the likely highly inflammatory environment produced by this coinfection exposure alters CD8^+^ T cell responses during leishmaniosis has not been explored. Unlike their CD4 counterparts, TBC+ dog CD8^+^CD49d^hi^ T cell proliferation and PD-1 expression in response to TLA-stimulation both moderately increased after the TBC seroconversion event (Figure 4C, D, and E) and were not significantly correlated considering all TBC+ timepoints (Figure 4F). At endpoint measurements, chronic TBC+ dog CD8^+^CD49d^hi^ T cell proliferation in response to TLA was significantly higher than in TBC-dogs, while acutely TBC+ dogs showed an intermediate level of proliferating cells (Figure 4L). CD8^+^CD49d^hi^ T cell proliferation did not significantly correlate with level of PD-1 expression at the endpoint measurements in any groups. Together this data indicates differential effects of TBC on antigen-induced proliferation of CD4^+^ vs CD8^+^CD49d^hi^ T cells. CD4^+^ T cell proliferation decreases after TBC seroconversion while CD8+ T cell proliferation increases.

To examine clinical diagnostic and TLA-stimulated immune readout relationships post-coinfection, Supplemental Figure 5 shows a correlation matrix and corresponding p-values, including the TBC seroconversion and subsequent timepoints. We wanted to establish whether the effect of increasing proliferating CD8 T cells was specific to the TBC+ dog group or applied to the entire cohort. Therefore, in Figure 5, we performed correlation analysis between proliferating CD4 or CD8 T cells and *Leishmania* parasitemia *vs*. serological results at the endpoint measurements including all subjects (Figure 5). Indeed, we observed that TLA-stimulated %CFSE^lo^ CD8^+^CD49d^hi^ T cells were positively correlated with *Leishmania* blood burden (Figure 5A, p=0.045) and *Leishmania* serological OD ratio (Figure 5B, p=0.015) but not LeishVet score. Enhanced proliferation among CD8^+^CD49d^hi^ T cells in this CanL cohort was associated with higher blood parasite burden.

**Figure 5.**
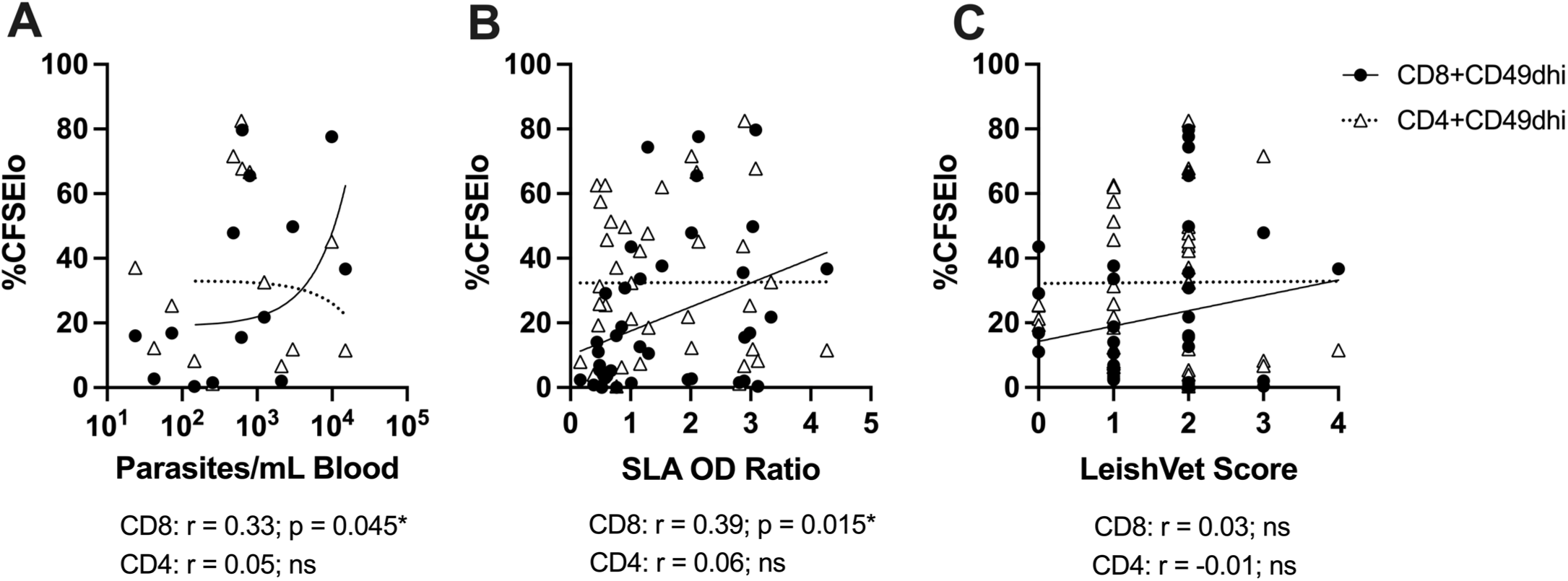
*Leishmania*-antigen stimulated CD8^+^ T cell proliferation significantly correlates with increasing *Leishmania* **burden and serology.** Correlation of endpoint measurement TLA-stimulated %CFSElo CD4^+^CD49d^hi^ (triangles, dotted line) or CD8^+^CD49d^hi^ (circles, solid line) lymphocytes and *Leishmania* diagnostic readouts including all study subjects. Spearman correlation coefficient (r), statistical result, and linear regression shown.

### Impact of coinfection on long-term clinical outcomes

Finally, we tested if our immune readouts could delineate dogs that developed severe leishmaniasis as measured by LeishVet status 3 or 4 or mortality due to leishmaniosis. After the conclusion of the immunological study period, remaining subjects were visited for CanL clinical evaluation. We stratified the cohort by mild or severe CanL at follow-up assessment and by TBC status during the study period (Figure 6). At the endpoint study measurement, dogs that developed severe CanL already displayed a signature of significantly increased blood parasite burden, anti-*Leishmania* serology, and LeishVet score compared to dogs maintaining mild disease (Figure 6A-C). Additionally, both antigen-stimulated IFNγ and, to a lesser extent, IL-10 secreted by PBMCs were already significantly reduced at the end of the study in dogs going on to develop severe CanL (Figure 6D-E).

**Figure 6.**
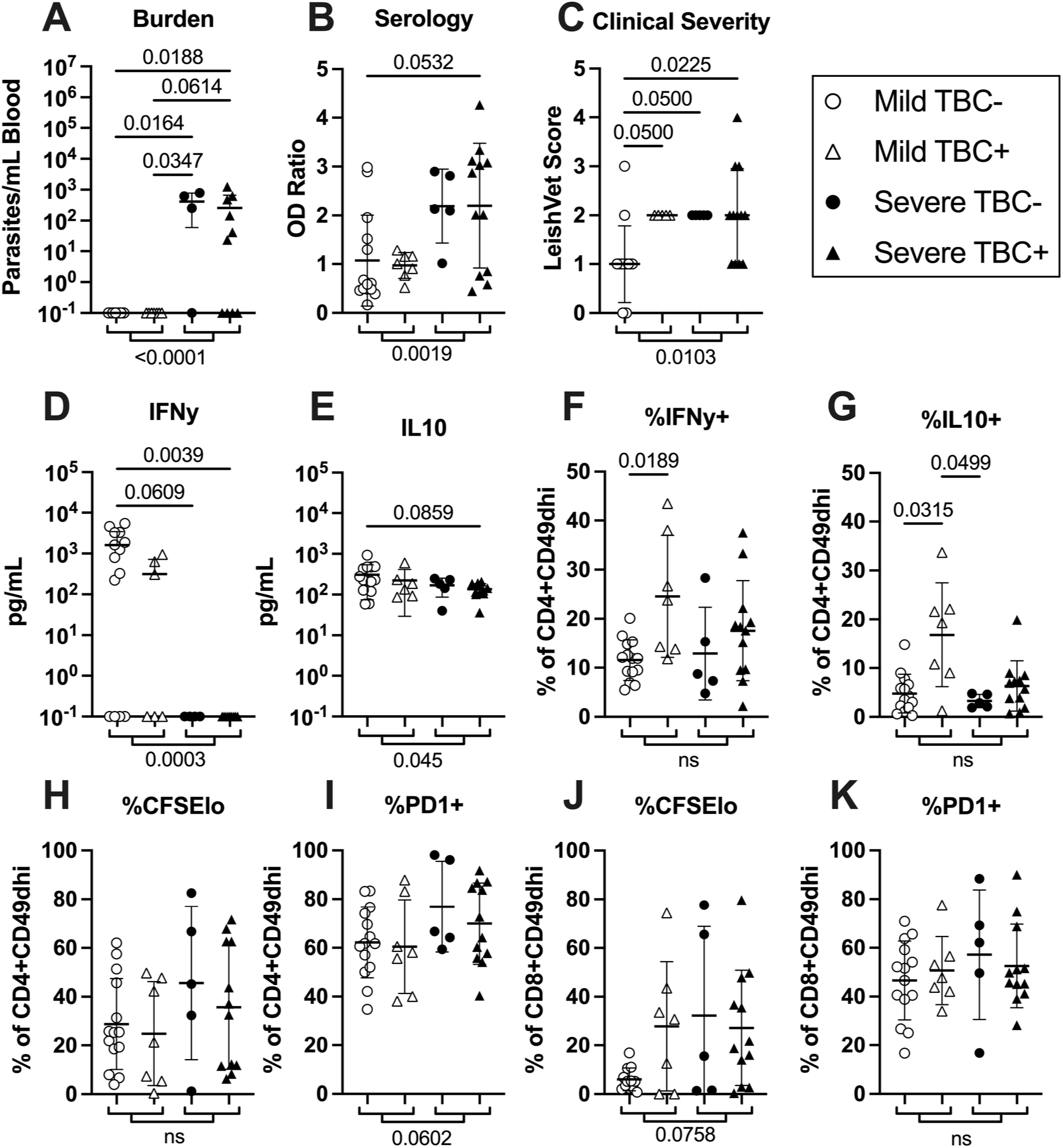
Decreased cytokine production and a trend of increased CD8^+^CD49d^hi^ T cell proliferation prior to development of severe CanL. Comparison of endpoint measurements of indicated readout among dogs that had developed severe CanL and/or were euthanized due to leishmaniasis at post-study follow up timepoints (Severe) vs dogs that displayed mild disease at follow up timepoints (Mild) separated by TBC status during the study period. Outliers removed. P-values denoted between individual groups: Kruskal-Wallis with Dunn’s post-test or One-Way ANOVA with Holm-Sidak post-test. P-values denoted between Mild and Severe groups: unpaired t-test or Mann-Whitney test. Mean and SD shown.

When splitting the clinical outcome groups into dogs which did or did not experience TBC during the main study period, TBC+ dogs that maintained mild CanL disease showed significantly higher proportions of both IFNγ^+^ and IL-10^+^ CD4^+^CD49d^hi^ T cells at the study culmination compared to TBC-dogs maintaining mild disease (Figure 6F-G). However, TBC+ dogs that went on to develop severe CanL did not show enhanced CD4^+^ T cell cytokine expression, perhaps indicating immunosuppression associated with the severe CanL may already be occurring during the study in this group.

The proportion of proliferating and PD-1 expressing T cells trended to be increased among dogs developing severe disease post-study (Figure 6H-K) compared to those maintaining mild CanL but did not show a significant difference driven by TBC status.

## DISCUSSION

Cross sectional studies showed dogs with CanL have a higher rate of coinfections with other vector-borne diseases than non-CanL dogs in *Leishmania* endemic areas (27–30). TBCs are highly prevalent canine coinfections in *Leishmania-*endemic countries due to the extensive distribution of tick vectors and range of associated tick-borne pathogens (15). Furthermore, TBCs have been associated with more severe clinicopathological abnormalities (*R. conorii, E. canis, A. phagocytophilum* and *B. henselae)* and higher LeishVet clinical scores (*A. phagocytophilum)* (29). Whether dogs with CanL are more susceptible to TBCs due to *Leishmania-*induced immunosuppression or if TBCs trigger evolution of underlying subclinical *Leishmania* infections, is not clear. This study is unique as we were able to follow the natural development of CanL and associated immune parameters, alongside concurrent TBC under real-world conditions.

Over the course of this 18-month study, subclinical *Leishmania*-exposed individuals had significant increases in parasitemia, anti-*Leishmania* serum antibodies, and associated progressive clinical disease. Within the study period, 50% of subjects developed serum antibodies reactive against TBC antigens. Secondary infection with TBC led to more severe clinical presentation compared to TBC-CanL dogs. In addition, dogs positive for TBC during the study showed a significantly increased risk of mortality or progression to severe CanL following the study conclusion compared to TBC-dogs, indicating TBC had long lasting impacts on the host. These results are consistent with findings from a previous prospective study associating TBCs with more severe CanL disease and mortality (13). In the TBC+ cohort, LeishVet score (based off complete blood count and serum chemistry abnormalities), *Leishmania* blood parasite burden, and *Leishmania* serological reactivity values all strongly significantly correlated with one another, supporting that these readouts together act as a practical gauge of disease severity in this cohort.

In CanL enzootic areas of Europe and Brazil, *E. canis* often predominates, with up to 80% of dogs seropositive for *E. canis* in previous coinfection studies in Brazil (13). In the current study, *Ehrlichia* spp. accounted for 63% of TBCs and the majority of long-term TBC. *Borrelia burgdorferi* accounted for 37% of TBCs and 6 out 7 *Borrelia* seropositive dogs were positive for ≤6 months (Supplemental Figure 1B). TBCs likely contribute to CanL progression via multifactorial processes. *Anaplasma* and *Ehrlichia* species are both intracellular bacteria that infect and replicate within phagocytes systemically, the same cell types parasitized by *Leishmania* (31,32). This significant overlap in affected tissues such as spleen, liver, and bone marrow may synergistically disrupt the normal function of these organs, accelerating clinical manifestations of disease. In a Brazilian study, dogs co-exposed to *L*. *infantum* and *E*. *canis* presented significantly more pronounced hematological and biochemical abnormalities including hypoalbuminemia, increased alkaline phosphatase, and lower hematocrit than dogs positive for *Leishmania* only (33). Intracellularly, tickborne pathogens employ multiple immune evasion mechanisms. For example, *Anaplasma* and *Ehrlichia* species have been shown to inhibit lysosomal fusion, downregulate MHC class II expression, decrease TLR and MAPK signaling, and prevent NADPH assembly (11). Pessôa-Pereira et al. found *in vitro* coinfection with *L. infantum* and *Borrelia burgdorferi* in a canine macrophage cell line resulted in significantly greater intracellular *Leishmania* infection rate and burden. This was associated with increased *SOD2* mRNA expression and decreased mitochondrial reactive oxygen species activity (34). We hypothesize TBCs facilitate *Leishmania* survival in part by hindering mechanisms controlling intracellular *Leishmania* replication.

Innate and adaptive cytokines induced by TBC may alter the effectiveness of the Type 1 immune response controlling *Leishmania* replication in dogs maintaining asymptomatic leishmaniosis. *Borrelia burgdorferi* infection induces a mix of Type 1 and Type 17 inflammatory as well as regulatory IL-10 responses from canine and human cells (34–38). *E. canis* induces IFNγ secreting cells in dogs (39). Both IFNγ and IL-17 producing T cells occurred during canine infection with *Ehrlichia chaffeensis* (40). In mice, CD4^+^ T cell-derived IFNγ was required for resistance against lethal *Ehrlichia* infection, while inflammatory CD8^+^ T cells contributed to pathogenesis (41,42). IL-10 is upregulated transiently during acute timepoints (43,44).

Together, these studies show TBCs can induce a mix of Type 1, Type 17, and IL-10 responses and other innate inflammatory cytokines may occur. We speculate the inflammatory milieu is altered by TBC systemically and at focal sites of bacterial infection, in dogs concurrently infected with *Leishmania* parasites. Skewing away from Type 1 mediator production, especially in an environment with ongoing IL-10 production such as CanL (8), could reduce the ability of IFNγ to activate macrophages to control *Leishmania* replication. Meanwhile, excess co-stimulation by innate inflammatory cytokines in settings of chronic antigen presentation, which occurs during CanL, is known to promote the onset of T cell exhaustion (25).

To uncover signs of potential immune exhaustion in coinfected dogs, we assayed T cell IFNγ and IL-10 production, proliferation, and PD-1 expression. We saw a transient increase in the amount of TLA-stimulated IFNγ produced by bulk PBMCs three months after TBC seroconversion, which declined progressively at six-and nine-months following TBC (Figure 3A). Similarly, CD4^+^CD49d^hi^ T cells showed an increased frequency of TLA-stimulated IFNγ production in dogs acutely TBC, but the increase subsided in dogs with chronic TBC exposure (Figure 3G). Transiently increased IFNγ production post-TBC may be driven in response to the bacterial infection directly in the form of a bacteria antigen specific cellular response, or in a bystander fashion due to Toll-Like Receptor activation or cytokines acting on existing effector T cells nonspecifically (45).

By 12 months post-TBC, CD4^+^CD49d^hi^ T cells showed enhanced IL-10 production in response to TLA (Figure 3G). Overall, we observed a trend of lower IFNγ/IL-10 production by CD4^+^ T cells in response to both *Leishmania* antigen and mitogen after a prolonged period of TBC compared to TBC-dogs. Other studies have observed lower or unchanged IFNγ output in response to *Leishmania* antigen during active VL or CanL using various antigen stimulation and molecular readout techniques (7,46–49). During active human VL, Leishmania antigen stimulated PBMCs produce IL-10 but low IFNy, which recovers after treatment (caldas).

Statistical modeling of subject-matched longitudinal data showed TBC is followed by moderate to strong probability of decreased CD4^+^ T cell proliferation in response to TLA (Figure 4). This was highly positively correlated with CD4^+^ T cell PD-1 expression in both TBC-and TBC+ subjects, indicating at this stage of CanL, PD-1 expression is serving as a marker of CD4^+^ T cell activation. The decrease in CD4^+^ T cell proliferation in our TBC cohort is not explained by a concomitant increase in PD-1 expression. Other studies have reported high CTLA-4 and PD-1 expression on CD4^+^ T cells during VL. While cells still produce IFNγ, blockade of inhibitory receptor-ligand interactions in infected macrophage co-cultures enhanced their ability to induce *Leishmania* killing (7,50), indicating these receptors are regulating functional capability. CD4^+^ T cell proliferation recovered after ∼9 months post-TBC and a difference was not observed at the endpoint compared to TBC-dogs. We did not assay the presence of CD4^+^ T cells responsive to tick pathogen antigens in this study.

In contrast to CD4^+^ T cells, TBC was associated with a moderate to strong probability that CD8^+^ T cells display increased proliferation and PD-1 expression in response to TLA following seroconversion, although these two factors were not significantly correlated (Figure 4). CD8^+^ T cell proliferation in response to TLA in coinfected dogs was significantly higher than seen in non-coinfected dogs at the endpoint measurements and was highest in dogs coinfected for longer periods of time (Figure 4L). Thus, TBC is progressively driving *Leishmania*-specific CD8^+^ T cells to proliferate over time.

We also saw that increased CD8^+^ T cell proliferation in response to *Leishmania* antigen correlated with increased parasite burdens and higher amounts of circulating anti-*Leishmania* antibodies (Figure 5), both associated with worsening disease. Cortese et al. previously found a significant increase in CD8^+^ lymphocyte frequency in dogs with CanL (51). Interestingly, *E*. *canis* infection in dogs has also been associated with a significant increase in CD8^+^T cells (52). Tickborne coinfections exacerbate CD8^+^ T cell proliferation in response to *Leishmania* antigen during CanL.

The roles of CD8^+^ T cells in human VL and CanL are not well studied compared to cutaneous leishmaniasis, where they contribute to both parasitic control and immunopathology through IFN-γ expression and cytotoxicity, respectively (53). It is thought that CD8^+^ T cells are more susceptible to immune exhaustion than their CD4^+^ counterparts, and CD8^+^ T cell dysfunction during VL has been previously reported (47,53–55). In humans with VL, circulating CD8^+^ T cells expressed significantly higher levels of IL-10 mRNA and CTLA-4 and PD-1 protein than endemic control cells, and these markers were even higher in splenic CD8^+^ T cells than PBMCs, where parasite load is concentrated (47). Blocking PD-1 ligand in CanL dog PBMC cultures lead to increased CD8^+^ T cell proliferation (7). VL patient CD8^+^ T cells appear activated with a CD45RO negative effector phenotype, expressing significantly higher levels of perforin and granulysin than endemic control cells, but did not contribute to *Leishmania* antigen-stimulated IFNγ (47,56). Additional inhibitory receptors LAG-3 and TIM-3 have also been found enriched on VL patient CD8^+^ T cells (57). Together, in VL, CD8^+^ T cells exist in an activated state with robust inhibitory receptor expression, potentially making them more susceptible to immune exhaustion. This is supported by our observation of TBC-driven exacerbated CD8^+^ T cell proliferation during CanL.

After the completion of this 18-month study, we assessed members of the cohort for mild or severe CanL and/or dogs that underwent owner-elected euthanasia due to leishmaniosis. Parasite burden, serological reactivity, and LeishVet score were significantly higher in dogs that had severe disease outcomes, while TLA-induced IFNγ and IL-10 production by bulk PBMCs were significantly reduced (Figure 6). However, the level of IL-10 remained quite high in both groups. Among cellular responses, both CD4^+^ T cell PD-1 surface expression and CD8^+^ T cell proliferation trended to be higher in dogs that later developed severe disease.

Tickborne coinfections promote clinical progression in dogs with asymptomatic CanL and have complex effects on the anti-*Leishmania* immune response, particularly leading to alterations in the IFNγ/IL-10 ratio and progressively increasing CD8^+^ T cell proliferation. More work needs to be done to explore the role of hyperactivation of CD8^+^ T cells and their dysfunction during VL. Preventing tick bites in subclinical *L. infantum*-infected individuals removes compounding effects of chronic exposure to TBC immune evasion mechanisms and inflammation that may contribute to the onset of immune exhaustion (58). Limiting the course of disease in dogs is important for reducing the risk of transmission to humans in endemic communities (10). Regular application of oral or topical acaricides, combined with environmental modifications, and manual tick checks are currently the most effective ways to prevent TBCs at present (59). Clinically, tick preventatives should be incorporated into the management of subclinical *L. infantum*-infected dogs to prevent TBCs and reduce the risk of progression to clinical leishmaniosis.

## ACKNOWLEDGEMENTS

A great amount of time, coordination, and teamwork was required to complete this study. We are forever thankful to the animal caretakers who participated in the study and worked with our team to organize site visits. We also express our deepest gratitude to all members of the Petersen lab and volunteers that assisted in site visits, sample collection, or sample processing. This research was supported by the MFHA Foundation and the University of Iowa Interdisciplinary Immunology Postdoctoral Training Grant NIH/NIAID T32AI007260.

